# Combining machine learning algorithms and single-cell data to study the pathogenesis of Alzheimer’s disease

**DOI:** 10.1101/2024.01.26.577320

**Authors:** Wei Cui, Liang Zhang, Fang-Rui Zheng, Xi Huang Li, Gui-Lin Xie

## Abstract

Extracting valuable insights from high-throughput biological data of Alzheimer’s disease to enhance understanding of its pathogenesis is becoming increasingly important. We engaged in a comprehensive collection and assessment of Alzheimer’s microarray datasets GSE5281 and GSE122063 and single-cell data from GSE157827 from the NCBI GEO database. The datasets were selected based on stringent screening criteria: a P-value of less than 0.05 and an absolute log fold change (|logFC|) greater than 1. Our methodology involved utilizing machine learning algorithms, efficiently identified characteristic genes. This was followed by an in-depth immune cell infiltration analysis of these genes, gene set enrichment analysis (GSEA) to elucidate differential pathways, and exploration of regulatory networks. Subsequently, we applied the Connectivity Map (cMap) approach for drug prediction and undertook single-cell expression analysis. The outcomes revealed that the top four characteristic genes, selected based on their accuracy, exhibited a profound correlation with the Alzheimer’s disease (AD) group in terms of immune infiltration levels and pathways. These genes also showed significant associations with multiple AD-related genes, enhancing the potential pathogenic mechanisms through regulatory network analysis and single-cell expression profiling. Identified three subpopulations of astrocytes in late-stage of AD Prefrontal cortex dataset. Discovering dysregulation of the expression of the AD disease-related pathway maf/nrf2 in these cell subpopulations Ultimately, we identified a potential therapeutic drug score, offering promising avenues for future Alzheimer’s disease treatment strategies.

## Introduction

Alzheimer’s disease (AD) is characterized by the dysfunction of several brain macroscale networks. At the subcellular level, this includes disturbances in amyloid protein-β (Aβ) and tau protein stability, inflammatory immune responses, lipid and bioenergy metabolism, oxidative stress, and other molecular mechanisms, as well as disorders in cell signal transduction pathways ^[1]^. Pathologically, AD manifests as tau-containing neurofibrillary tangles, Aβ plaques, neuronal loss, glial hyperplasia, or enlarged endosomes ^[2]^. The primary role of proteins related to the amyloid protein cascade hypothesis is central in the pathogenesis of Alzheimer’s disease ^[3]^. The rapid advancement of microarray and high-throughput sequencing technology provides a robust and expansive approach for decoding the genetic and epigenetic underpinnings of diseases. Concurrently, this technology offers a wealth of evidence for the diagnosis and treatment of various conditions ^[4]^. With the progress of high-throughput sequencing technology and the proliferation of omics research involving large sample sizes, the AD regulatory network is becoming increasingly complex ^[5]^. Therefore, there is an urgent need for more precise diagnosis and treatment methods for AD. Traditional statistical methods are inadequate for the analysis of extremely high-dimensional and sparse omics data, leading to the rise of machine learning as the preferred method for complex data analysis ^[6]^. Leveraging machine learning to explore the pathogenesis and drug response mechanisms of complex diseases based on multi-omics data will significantly advance precision and translational medicine ^[7] [8]^.

In our research, we combined two datasets and identified 148 differentially expressed genes between the control group and the AD group using the limma package. GO analysis and KEGG pathway enrichment analysis of these differentially expressed genes revealed their close relationship with the pathogenesis of AD. For feature selection of DGEs, we applied Lasso regression and the SVM algorithm, and the SVM-RFE algorithm was utilized to identify the top four feature genes with the highest accuracy. Focusing on these key genes, we conducted analyses of immune cell infiltration levels, enrichment pathways, tSNE algorithm-based single-cell expression distribution, expression of key genes in subtype pathways, and cMap drug prediction. These analyses provided insights into the potential of key genes as targets and illuminated the potential molecular mechanisms that influence disease progression.

## Results

### 1. Screening of DEGs

We sourced the GSE5281, GSE122063, and other Alzheimer’s disease-related datasets from the GEO database, encompassing a total of 261 patient expression profile datasets, which included 118 normal samples and 143 disease patient samples. The Combat algorithm was employed for chip correction, and the PCA plot was utilized to illustrate the differences pre- and post-correction. Post-correction results indicated a reduced batch effect between chips, as demonstrated by the Combat algorithm (Fig. 1.A-Fig. 1.B). Subsequently, the limma package was applied to determine differentially expressed genes between the two patient groups, adhering to the screening criteria of P.Value < 0.05 & |logFC| > 1. This process identified a total of 148 differentially expressed genes, comprising 76 up-regulated and 72 down-regulated genes (Fig. 1.C). Further, we conducted pathway analysis on these 148 differentially expressed genes. The results revealed that these genes were primarily enriched in pathways such as cell body, brain development, response to steroid hormone, and embryonic morphogenesis, among others (Fig. 1D-F).

**Figure 1.**
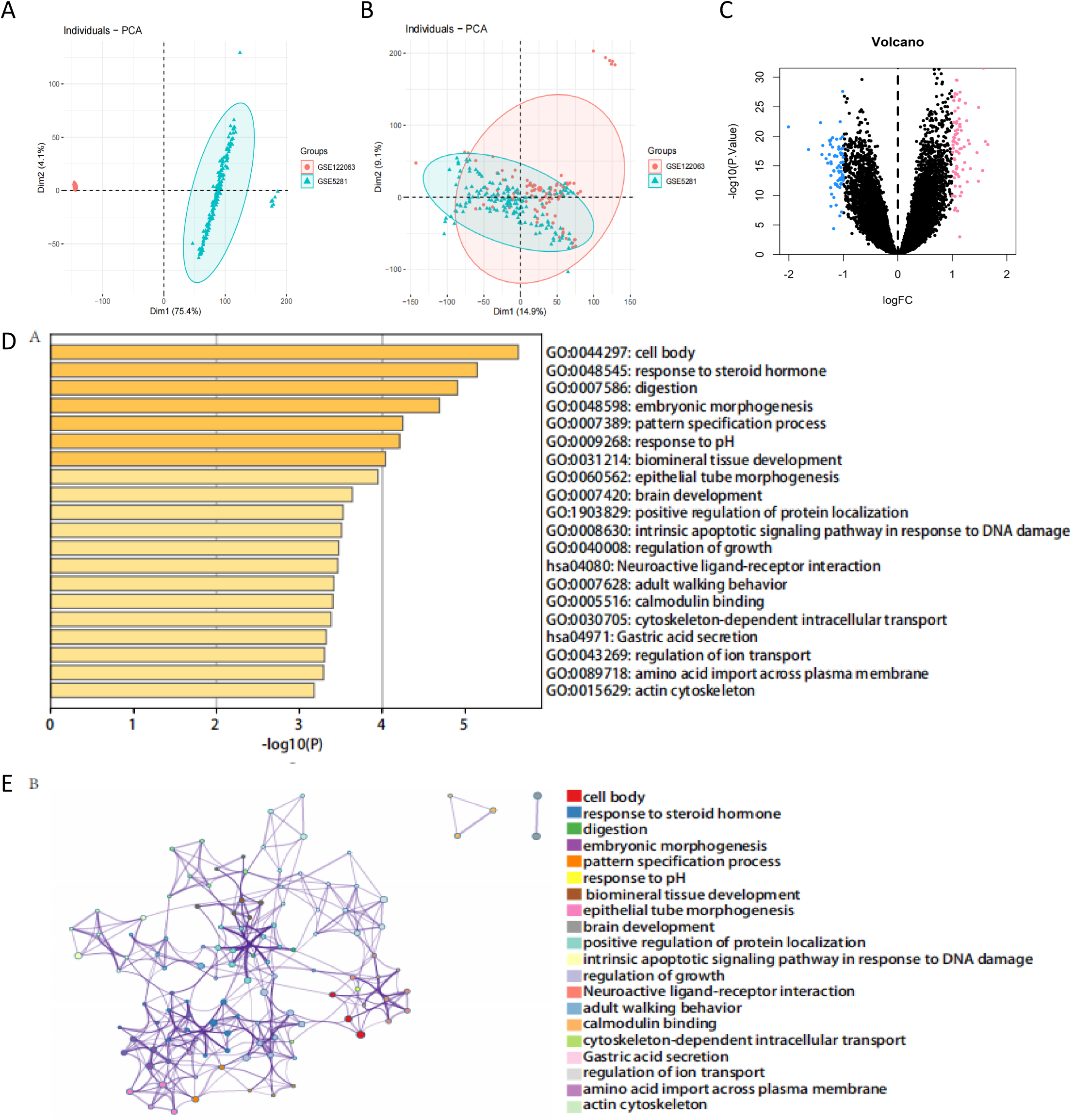
Identification of differentially expressed genes. **A.** PCA plot of disease group. **B.** PCA plot of normal group (red) and disease group (green). **C.** Volcano plot of DEGs between disease group and normal group (green: down-regulated DEGs; red: up-regulated DEGs) P<0.05, |log2FC|>2. D.Functional analysis of differentially expressed genes between disease group and normal group **D.** GO enrichment results of differentially expressed genes between the disease group and the normal group (p value<0.01) **E.** Protein interaction network of differential genes.

### 2. Functional enrichment analysis and identifiation of key genes

To further pinpoint key genes impacting Alzheimer’s disease, we concurrently employed lasso regression and SVM feature selection algorithms to filter through 148 differentially expressed genes, identifying characteristic genes associated with Alzheimer’s disease. Lasso regression pinpointed a total of 42 genes as characteristic of Alzheimer’s disease (Fig. 2.A-Fig. 2.B). Simultaneously, we employed the SVM-RFE algorithm to evaluate characteristic genes in Alzheimer’s disease. This approach, focusing on the top four characteristic genes with the highest accuracy, yielded four intersection genes derived from the set of genes identified by the lasso regression algorithm (Fig. 2.C-Fig. 2.D). These four genes, CDC37, LOC100272216, MAFF, and MYL5, are designated as key genes for our subsequent research.

**Figure 2.**
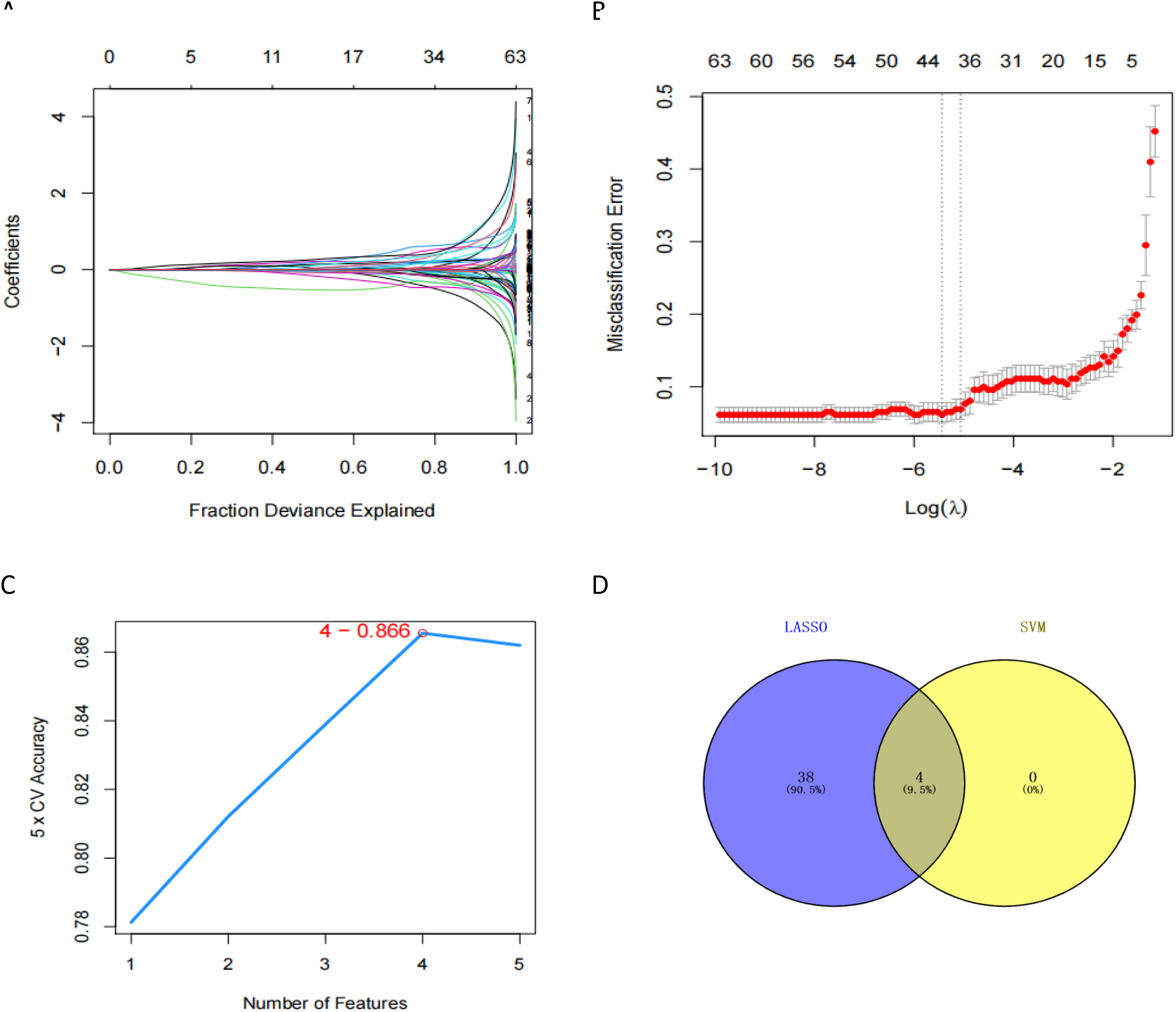
Selection of diagnostic biomarkers and identification of hub genes. **A.** LASSO analysis cross-validation identified 42 genes. **B.** LASSO coefficient spectrum of differentially expressed genes. **C.** SVM-RFE prediction of true value change curve. **D.** Venn diagram of Lasso screening results and SVM-REF screening results.

### 3. Estimating of the relationship between hub genes and immune infiltration

The immune microenvironment is mainly composed of immune-related fibroblasts, immune cells, extracellular matrix, various growth factors, inflammatory factors, and specific physical and chemical characteristics. The immune microenvironment significantly affects disease diagnosis, survival outcome, and clinical treatment sensitivity. By analyzing the relationship between key genes and immune infiltration in disease datasets, we explored the mechanism by which key genes affect the progression of Alzheimer’s disease. The content of immune cells in each patient is shown in Fig. 3.A. There are multiple significant correlations between the levels of immune infiltration (Fig. 3.B). Compared with normal patients, the levels of Macrophages M1 and Macrophages M2 in disease group samples are significantly higher (Fig. 3.C). We further explored the relationship between key genes and immune cells, and found that multiple key genes are highly correlated with immune cells. CDC37 is positively correlated with NK cells activated, T cells follicular helper, and negatively correlated with Mast cells resting, B cells naive, etc. (Fig. 3.D); LOC100272216 is positively correlated with Macrophages M1, Plasma cells, and negatively correlated with Eosinophils, Dendritic cells activated, etc. (Fig. 3.E); MAFF is positively correlated with Neutrophils, B cells naive, and negatively correlated with NK cells activated, B cells memory, etc. (Fig. 3.F); MYL5 is positively correlated with B cells memory, T cells CD8, and negatively correlated with T cells CD4 memory resting, B cells naive, etc. (Fig. 3.G). Then, we obtained the correlation between these key genes and different immune factors from the TISIDB database, including immune inhibitors, chemokin es, immune stimulators, receptors, and MHC (Fig.3.H). These analyses confirm that these key genes are closely related to the level of immune cell infiltration and play an important role in the immune microenvironment.

**Figure 3.**
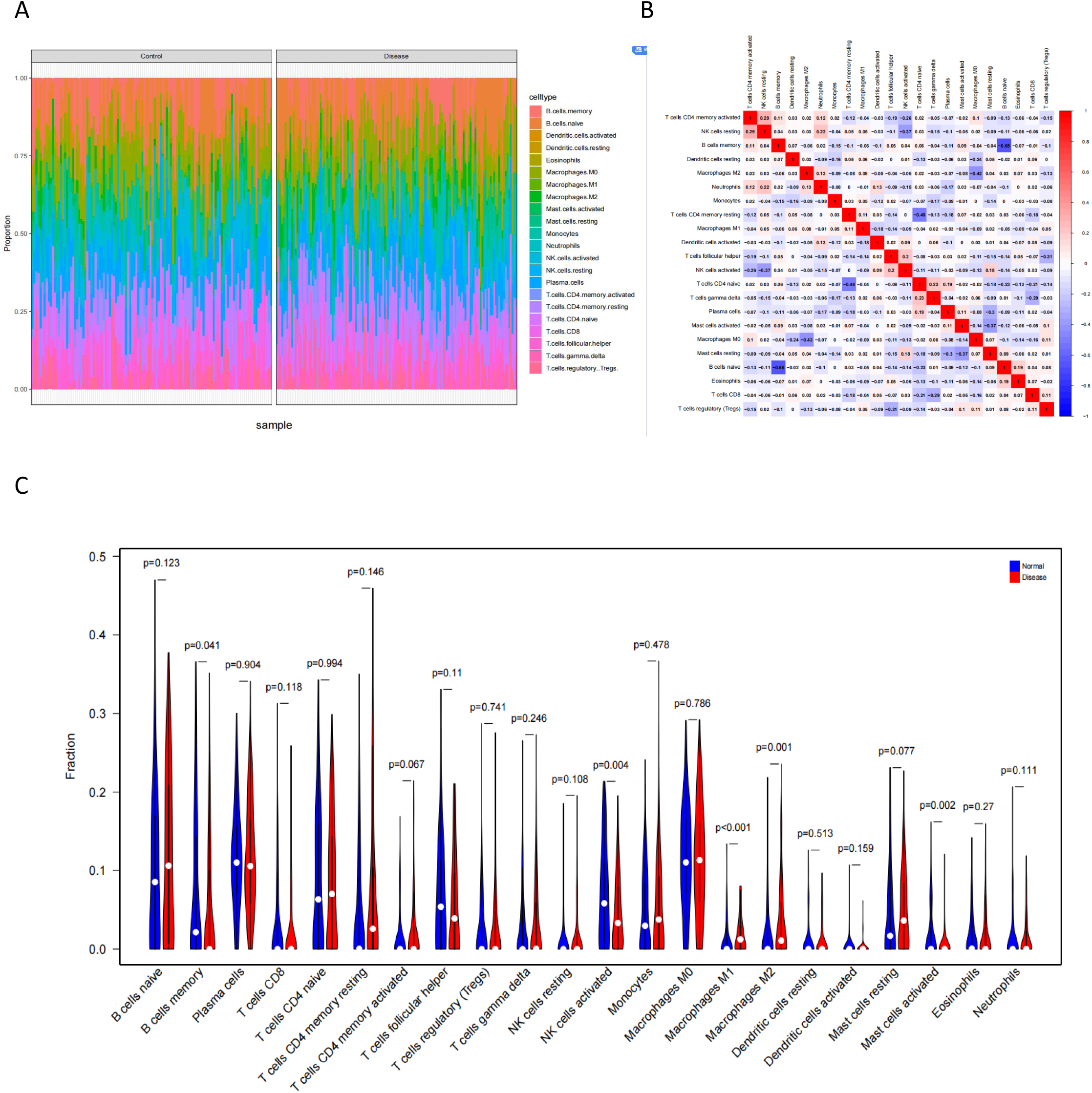
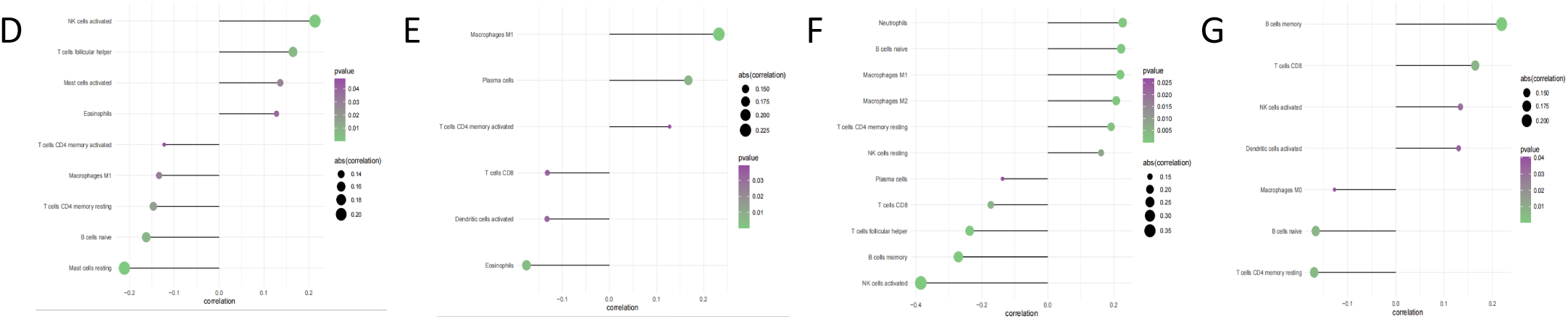
Immune infiltration between AD and normal controls. **A.** Cumulative histogram of the relative percentage of infiltrating immune cells in the disease group and normal group. **B.** Correlation between immune cells. **C.** Differences in immune cells between the disease group and normal group (P<0.05 is statistically significant) (red is the disease group, blue is the normal group) **D-G.** CIBERSORT’s correlation with the expression of CDC37, LOC100272216, MAFF, and MYL5, respectively.

### 4. Identification of the potential mechanisms of the key genes by GSEA

We subsequently delved into the specific signaling pathways associated with key genes, aiming to elucidate the potential molecular mechanisms through which these genes influence disease progression. The results of the Gene Set Enrichment Analysis (GSEA) revealed distinct pathways enriched by the high expression of each key gene. Specifically, high expression of CDC37 was predominantly enriched in pathways such as KEGG Parkinson’s Disease(Supplementary materials, S.1), KEGG Huntington’sDisease(S.1), and KEGG Oxidative Phosphorylation (Fig.4.A), among others. In the case of LOC100272216, its high expression was mainly enriched in pathways like KEGG NOTCH Signaling Pathway(S.1), KEGG Adhesion Junction(S.1), and KEGG Chronic Myeloid Leukemia(S.1). For MAFF, the significant enrichment was observed in pathways including KEGG Chronic Myeloid Leukemia, KEGG NOTCH Signaling Pathway(Fig.4.B), and KEGG Small Cell Lung Cancer. Lastly, the high expression of MYL5 was mainly enriched in pathways such as KEGG Nicotinate and Nicotinamide Metabolism (Fig.4.D), KEGG Aminoacyl-tRNA Biosynthesis (Fig.4.C), and KEGG Alanine, Aspartate, and Glutamate Metabolism, among others (S.1). These findings indicate that the high expression of these key genes is intricately linked to various signaling pathways, which potentially play a crucial role in the progression of Alzheimer’s disease and related disorders.

**Figure 4.**
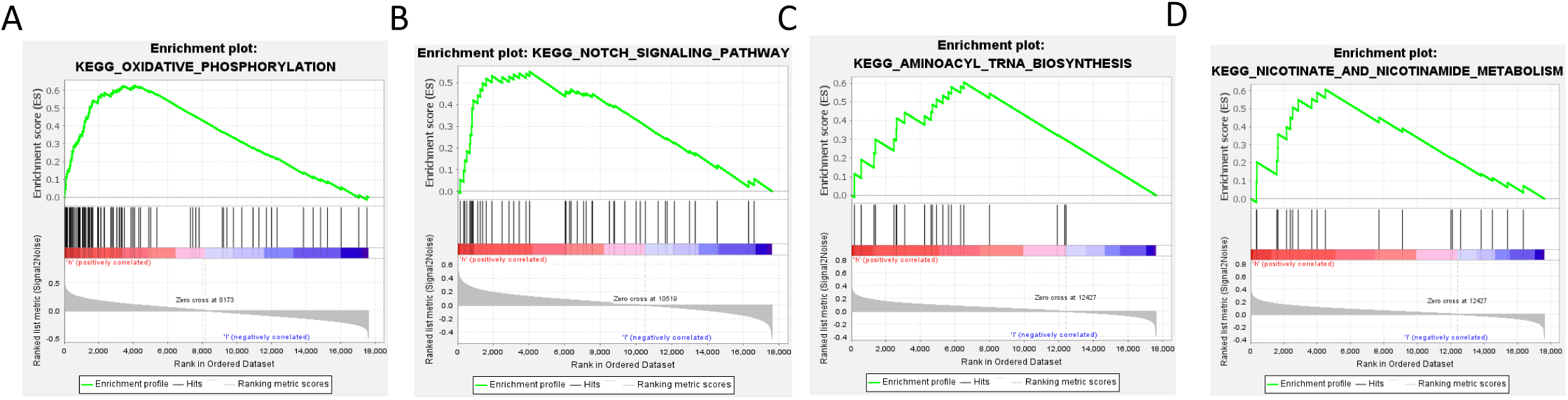
GSEA enrichment pathways of key genes. **A-D.** CDC37, MAFF, MYL5, and MYL5 are respectively enriched in these specific pathways.

### 5. Enrichment analysis for transcription factors of key genes

In this analysis, the four key genes were included in a gene set, revealing that they are regulated by multiple transcription factors and other common mechanisms. Consequently, an enrichment analysis was conducted, focusing on these transcription factors. This involved the use of cumulative recovery curves (Fig.5.A-B), Motif-TF annotation, and the selection and analysis of important genes. The analysis results indicated that the motif with the highest Normalized Enrichment Score (NES: 5.59) was cisbp M1140. This particular motif was found to be enriched in two of the key genes, specifically MAFF and MYL5. Subsequently, a comprehensive display of all enriched motifs and their corresponding transcription factors for these key genes was presented (Fig.5.C-E). These findings underscore the intricate regulatory network surrounding these key genes, highlighting the significance of specific motifs and transcription factors in their regulation. This insight is instrumental in understanding the molecular mechanisms through which these genes potentially influence the progression of Alzheimer’s disease and other related conditions.

**Figure 5.**
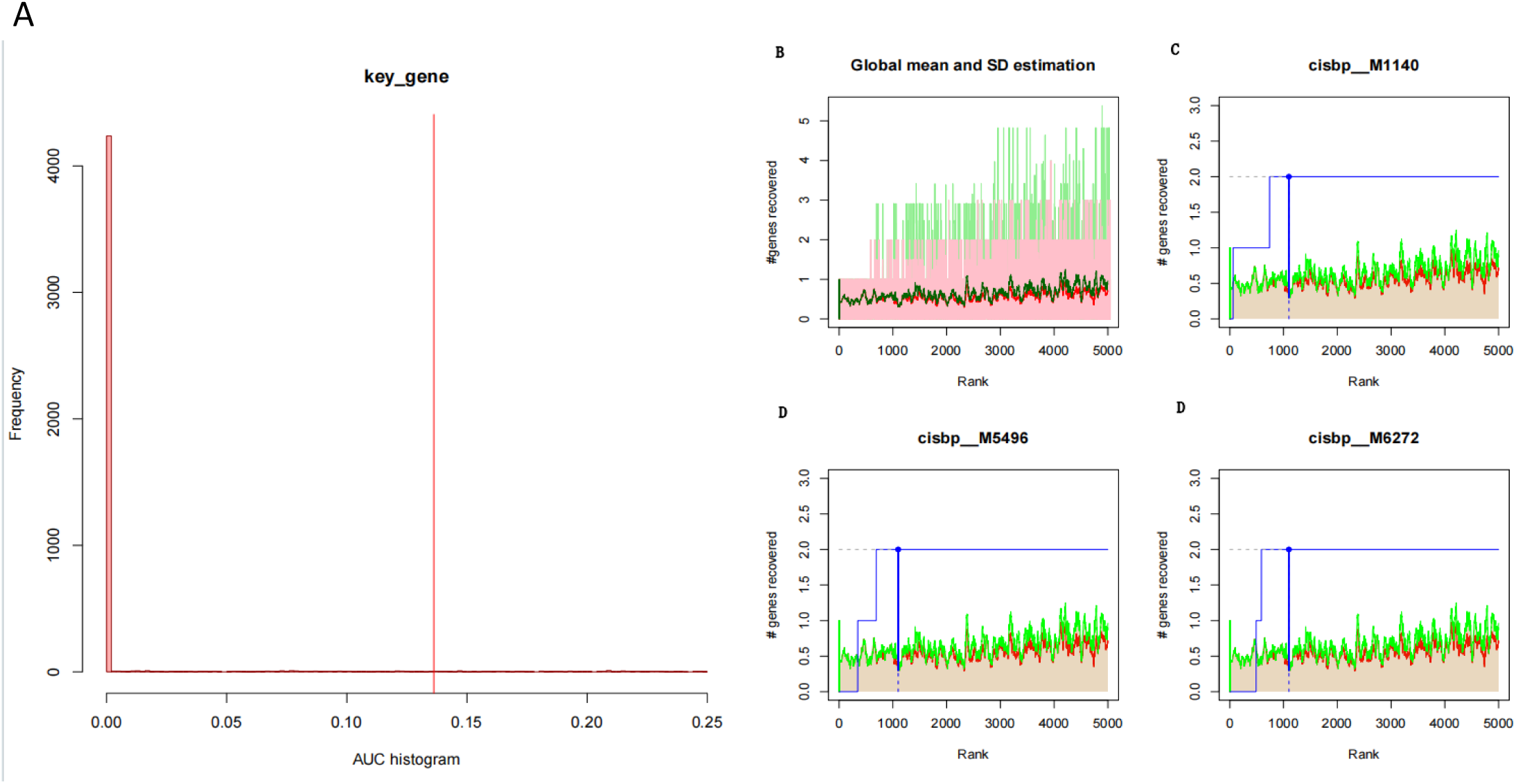
Transcription factor enrichment analysis (motif enrichment). **A:** Histogram of AUC. The overexpression of each motif in four key genes was evaluated by calculating AUC. The red vertical line indicates the significance level, and motifs with AUC higher than the significance level are considered significant motifs. **B:** Three important motif recovery curves. The red line represents the global average of the recovery motif curve, and the green line represents the average ±standard deviation. Motifs greater than the average ±standard deviation are considered statistically significant. The blue line represents the recovery curve. **C-E.**a display of enriched motifs and their corresponding transcription factors for these key genes was presented

### 6. Analysis of the relationship between key genes and the disease genes

Utilizing the GeneCards database (https://www.genecards.org/), we examined the expression differences of Alzheimer’s disease-related pathogenic genes between two groups. This analysis revealed distinct expression levels of several genes, including HFE, LMNA, LRRK2, MAPT, MPO, PSEN1, PSEN2, and SNCA, between the groups (Fig. 6.A). Furthermore, we carried out a correlation analysis between key genes and Alzheimer’s disease regulatory genes. The expression levels of key genes showed significant correlations with multiple Alzheimer’s disease-related genes. Notably, MAFF exhibited a substantial negative correlation with SNCA (Pearson r = -0.49), and CDC37 displayed a significant positive correlation with SNCA (Pearson r = 0.6) (Fig. 6.B).

**Figure 6.**
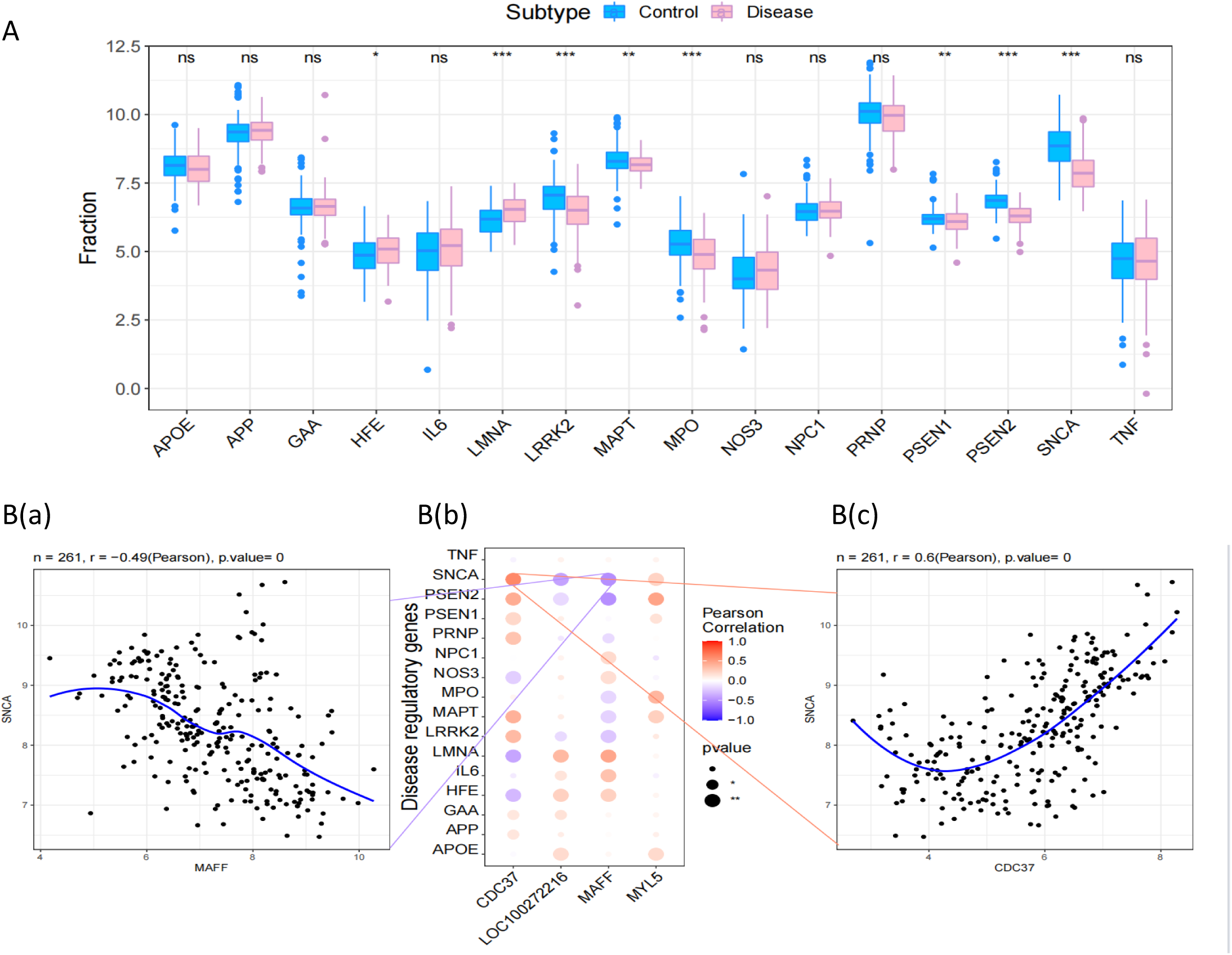
Correlation between ad-regulated genes and core genes. **A:** Ad regulates the differential expression of genes. (\**p* < 0.05 is considered to be a significant difference \*\**p* < 0.01, \*\*\**p* < 0.001, ns, no significant) **B:** Pearson correlation between key genes and differentially regulated genes. Blue indicates negative correlation, and red indicates positive correlation. **a.** Maff is significantly positively correlated with SNCA. **b.** Visualized Pearson correlation between hub genes and differentially regulated genes. The graph shows a significant positive correlation between CDC37 and SNCA. The Pearson coefficient and p-value are displayed at the top of the graph (*p* < 0.01 indicates a significant correlation).

### 7. Expression of key genes in scRNA-seq and Potential drugs prediction by connectivity map

Moreover, we utilized the Connectivity Map database to predict drugs for differentially up- and down-regulated genes. The results indicated that the expression profiles of drug perturbations from Rucaparib, Lestaurtinib, TCS-359, and Alvocidib were significantly negatively correlated with disease perturbation expression profiles. This suggests that these drugs could potentially alleviate or even reverse the disease state (Fig. 8.A-D).Finally, we analyzed the expression of key genes at the single-cell level. The expression of these key genes in Astrocytes, Macrophages, and Endothelial cells is depicted in the figure (Fig. 8.E-F). These findings offer valuable insights into the genetic underpinnings of Alzheimer’s disease, paving the way for targeted therapeutic strategies and furthering our understanding of the disease’s molecular landscape.

**Figure 8.**
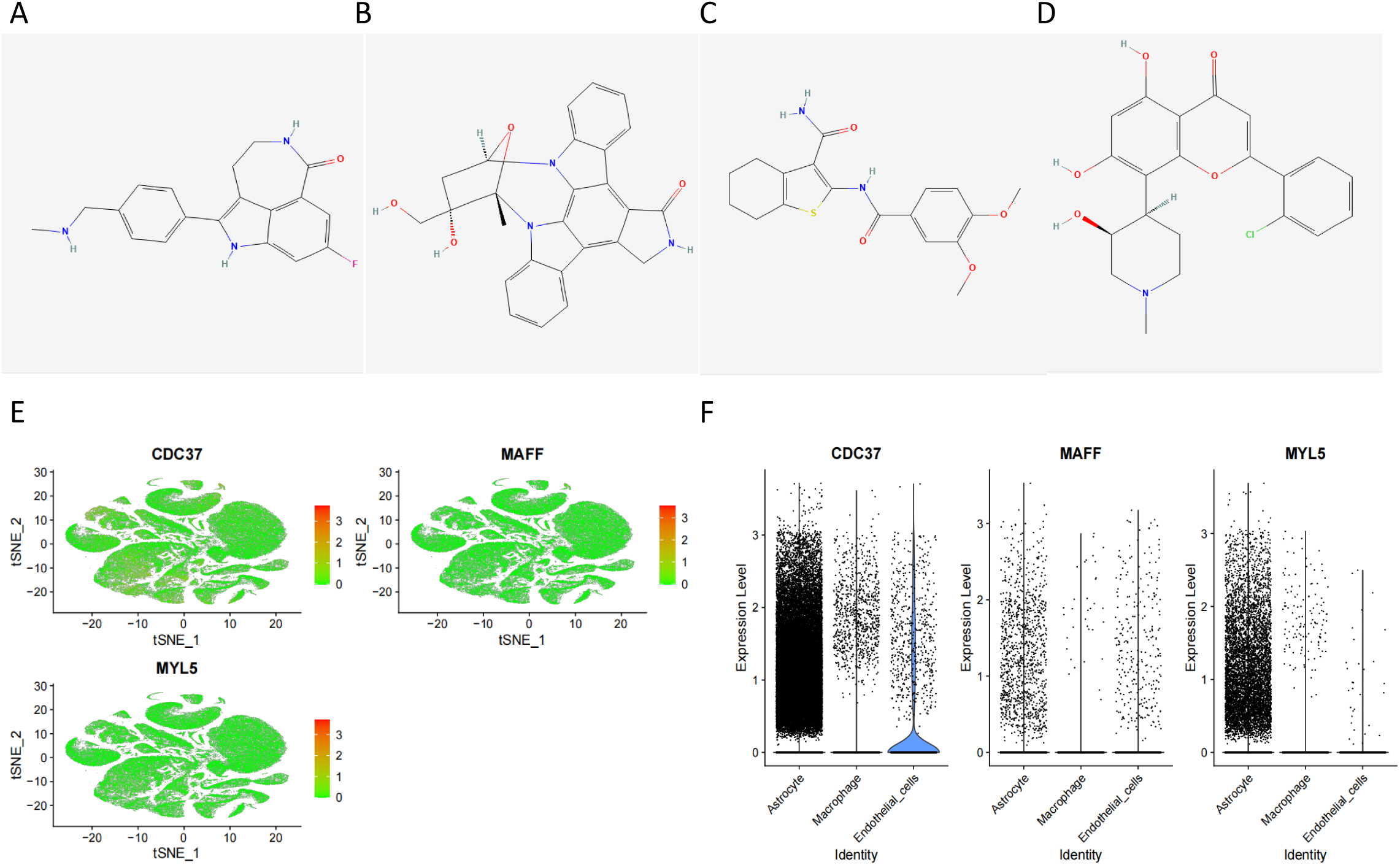
Drug prediction using cMAP A-D. Rucaparib, Lestaurtin ib, TCS-359, Alvocidib. **D:** Expression of core genes in five cell subpopulations. Three core genes are highly expressed in monocytes ; E: Violin plot of the expression of 3 core genes in 5 cell subpopulation.

### 8. Transcriptome specific astrocytes in the prefrontal cortex

In Braak staging late stage AD prefrontal cortex single-cell transcriptome dataset, the initial dataset consists of 20993 nuclei, of which 20186 nuclei passed QC and bimodal removal. Among them, 1972 were identified as astrocytes (Fig9. A). Three cell subpopulations with unique transcriptomic characteristics were identified by nuclear reprogramming analysis (Recluster) of astrocyte like expression (Fig9. B). To measure the activation level of Nrf2 Pathway in these cell subpopulations, AUCell was used for single-cell gene set evaluation. Therefore, we focused on subpopulation 0 as our research focus. MAFF with the highest variation in subpopulation 0 (Fig9. E).

We calculated the proportion of each subpopulation in the normal AD group and determined the largest cluster 0 (S2 supplementary material), which has different characteristics amd distribution from the previously reported astrocyte subpopulation [71]including high expression of SLC1A3 (cerebrusal fluid glucose transporter), SLC1A2, and CLU (Cluster Cell Protein) (Figure 9.C). They were evaluated through GO/pathway analysis to infer potential biological correlations (Fig. 9. D. Supplementary material S2). The MAPK Pathway (Mitogen Activated Protein Kinase), JAK-STAT Pathway (Janus Kinase Signal Producer and Activator of Transcription), and ECM Receiver Interaction Pathway in astrocyte cluster 0 mainly involve unique transcripts of intracellular and extracellular signaling, cell proliferation, differentiation, and survival, apoptosis, immune function, development, and repair. Clusters 1 and 2 of astrocytes do not significantly express the above pathways, but unlike the highly expressed functional pathways of cluster 1, they mainly express ribosome, mineral absorption, and histidine metabolism pathways, which are enriched in synthesis and absorption pathways. In order to investigate the effect of MAFF as a transcription factor at the subcellular level of the nrf2 pathway, we subsequently investigated the single-cell gene set score of the nrf2 regulatory pathway. Multiple downstream regulatory genes of nrf2 were highly expressed in subpopulation 0(Fig.9.E),, while multiple genes were not expressed in subcellular cluster 0 at the subcellular level(Fig.9.F).

**Figure 9.**
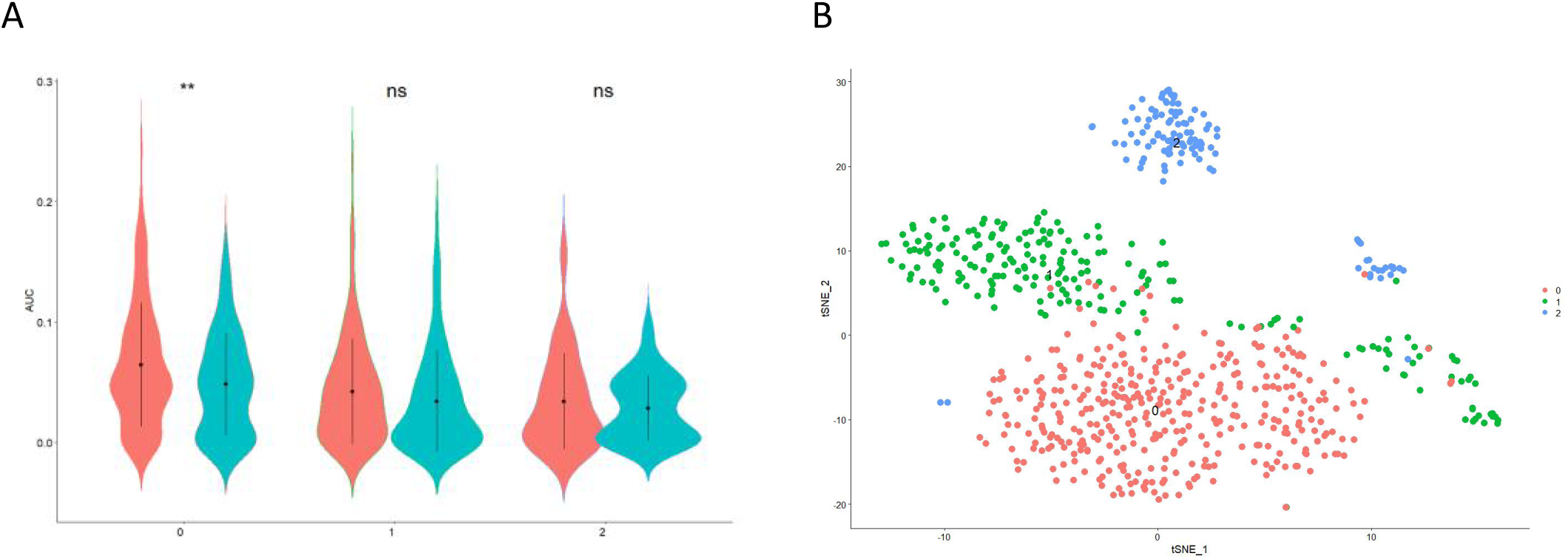
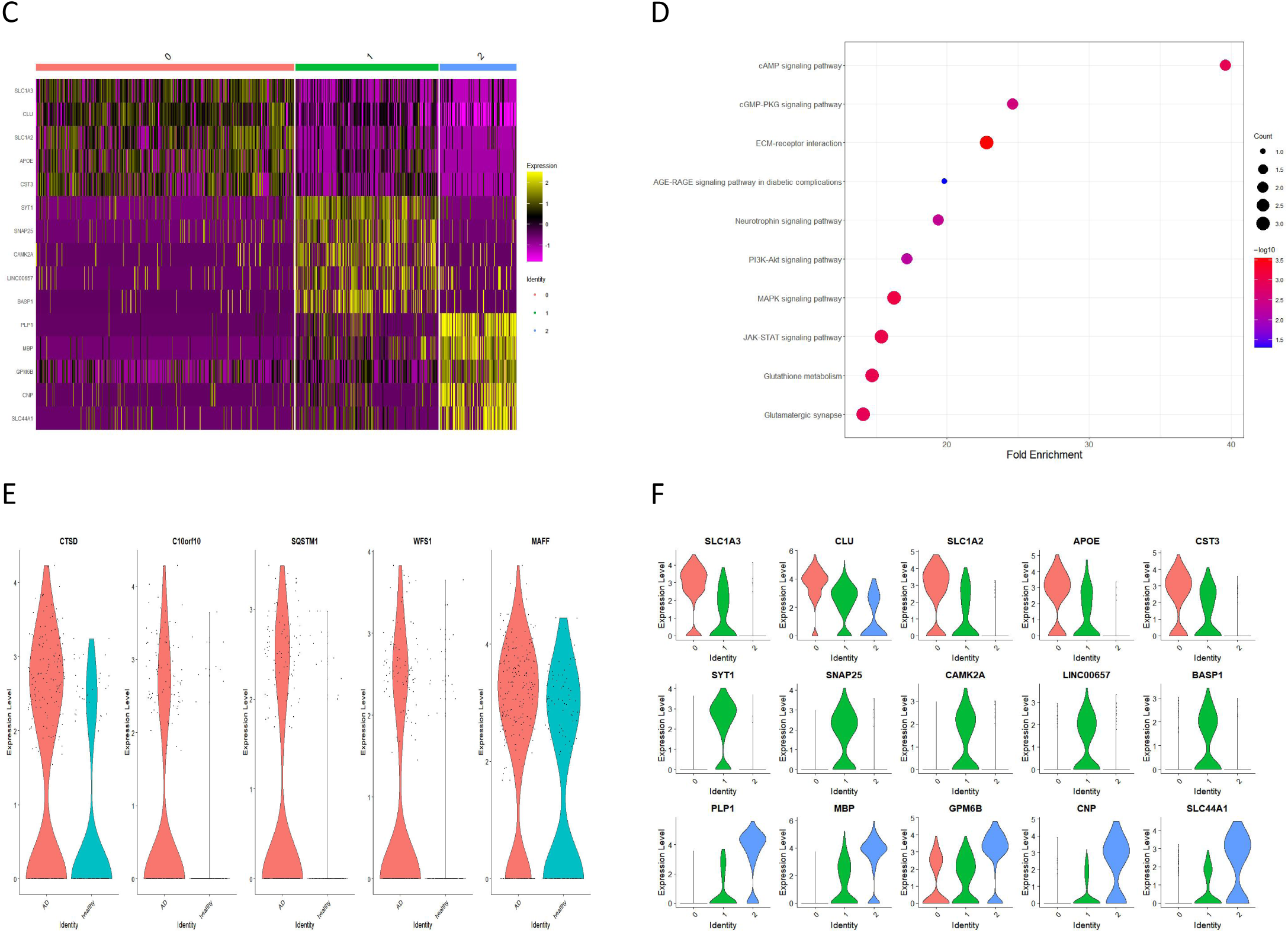
The state of astrocytes has different gene expressions and is related to biological pathways. **A)** Violin plot of AUC scores for different cell subpopulations **B)** TSNE graph after further grouping **C)** The expression heatmap of the top 5 marker genes in each cluster. **D)** KEGG analysis showed that each cell subpopulation exhibited different biological pathways. **E)** The gene with the greatest difference in expression between the AD group and the healthy group in subpopulation 0 **F)** Violin diagram of the expression of the first 5 marker genes in each cluster

**Figure 10.**
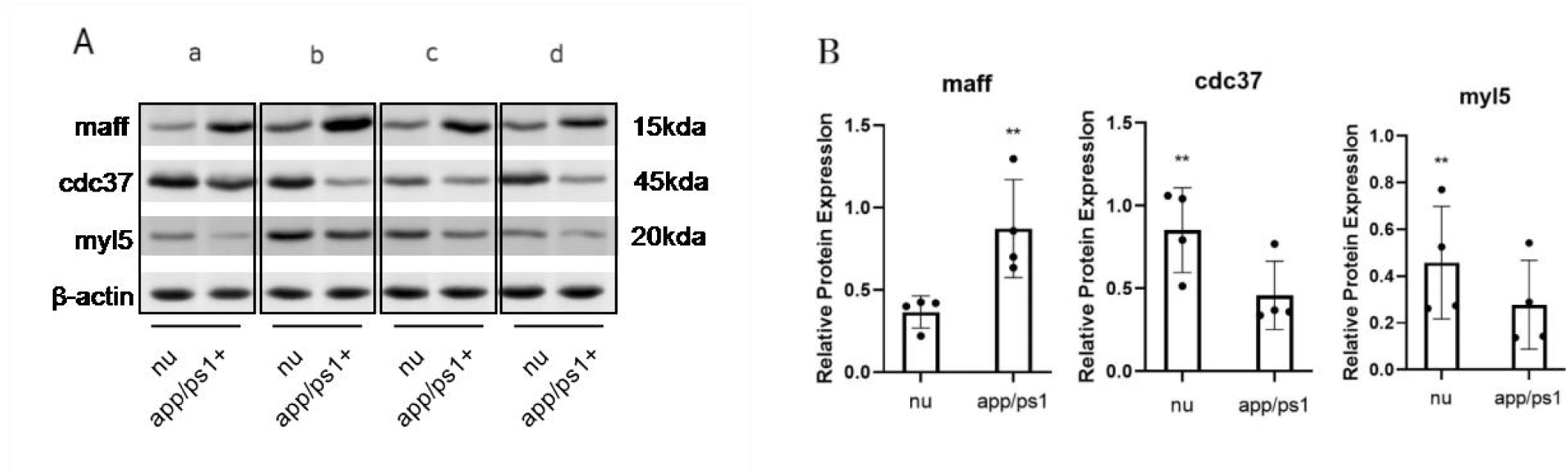
A: The WB expression levels of MAFF, CDC37, and MYL5 in the brain homogenate extract of App/Ps1 transgenic mice (n=3). B : Quantitative chart(\**p* < 0.05 is considered to be a significant difference \*\**p* < 0.01, \*\*\**p* < 0.001, ns, no significant)

### 9. Verification of key genes through Western blotting (WB)

APP/PSEN1 is a mature and widely used method for preclinical modeling of AD. We use WB to detect the expression levels of MAFF, CDC37, and MYL5 proteins. As shown in the figure, In the four repeated experimental groups, the expression of MAFF was significantly upregulated in the AD mouse group. CDC37 and MYL5 were significantly downregulated in model mice.

## Discussion

In this study, Our innovative approach of combining SVM RFE and Lasso algorithms with single-cell analysis used for AD research. The evaluation of characteristic genes showed a high accuracy rate (0.8 -0.866). Lasso regression, a widely-used machine learning model[9]. The accuracy of the Lasso model was ascertained by plotting the ROC curve. SVM-RFE, a feature selection algorithm based on the support vector machine (SVM)[10].

The expression levels of the top four characteristic genes with the highest accuracy were found to be strongly correlated with multiple genes related to molecular mechanism of AD and the immune infiltration level of AD.

Multiple pathways related to key genes have a significant impact on AD. The NOTCH signaling pathway, which have been proved that may play a key role in the vascular and neurodegenerative forms of cognitive impairment and dementia[11]. In the context of AD, MAFF can indirectly inhibit various detoxification and metallothionein genes [29–31]. This imbalance in signaling pathways may exacerbate oxidative stress [32] and disrupt synaptic plasticity [33, 34]. CDC37, as a co-chaperone protein, has been shown to regulate the maturation of kinases in the OXIDATIVE PHOSPHORYLATION pathway, [12] Cell Division Cycle Protein 37 (CDC37), in conjunction with HSP90, is instrumental in maintaining the stability of myelin formation and axonal integrity [22–25]. CDC37 has been identified as a significant hub in protein interaction analyses involving astrocytes exposed to stress or inflammation-inducing molecules [50].MYL5, the regulatory light chain 5 of the Myosin Light Chain (MLC). [44]. Evidence also indicates that the RhoA signaling pathway, which involves β-amyloid aggregation, tau phosphorylation, neuroinflammation, and synaptic damage, plays a significant role [46], MYL5 being key downstream effectors of RhoA signaling.

Multiple studies have shown that MAFF is significantly upregulated in AD under oxidative stress conditions, The transcription factor Nrf2, predominantly located in astrocytes, protects against pathological cell growth, including in degenerative diseases. MAFF regulates Nrf2 activity through the availability of these small MAF proteins [51] [52].MAFF is an essential co transcription factor in the transcription of the gene cascade containing antioxidant response elements (ARE) in the promoter region activated by heterodimerization between Nrf2 and one of the sMaf (myofascial fibrosarcoma oncogene homologs) proteins [70]. In the context of AD, oxidative stress is a significant factor, contributing to neuronal damage and the progression of the disease. The Nrf2 pathway’s activation enhances cellular defense mechanisms against oxidative stress, suggesting its potential therapeutic importance in conditions like AD [68]. Analysis of a large number of AD datasets has identified dysregulation of the NRF2 regulatory network, with a significant negative correlation between downregulation of genes with Nrf2 binding antioxidant response elements (ARE) and MAFF expression levels, which may play a role in this dysregulated network[72][73].

We found that in the late stage of AD Prefrontal cortex dataset, a subtype of astrocytes cells dominates. The differences in activation pathways, marker genes and types between astrocyte subpopulations and mixed subpopulations in Braak VI stage AD patients may indicate that the progression of AD cell subpopulations varies at different stages[71], They exhibit unique gene expression profiles and play a unique role in the development and severity of the disease. By analyzing the Nrf2 pathway, many downstream ARE element activation genes are not expressed in this subtype, but are highly expressed in other types of glial cells, indicating that the dysregulated genes of these nrf2 pathways may be differentially regulated in astrocytes. Further research is needed to investigate the differential regulatory network dysregulation of transcription factors MAFF and Nrf2 pathways in astrocyte AD subpopulations.

In conclusion, the comprehensive analysis of the AD dataset from GEO suggests that CDC37, MYL5, MAFF, and LOC100272216 are promising therapeutic targets. The application of machine learning algorithms, including RF and SVM-RFE, demonstrates robust predictive performance, paving the way for novel treatment strategies for AD patients.

## Materials

### 1. Data Acquisition

We procured the Series Matrix File of GSE5281 from the NCBI GEO public database, complemented by the annotation file GPL570. This dataset comprises expression profile data from 161 samples, including 74 normal samples and 87 patients with the disease. Additionally, we downloaded the Series Matrix File of GSE122063, annotated with GPL16699, encompassing expression data from 100 samples, with 44 normal and 56 disease patients. These datasets were amalgamated, and batch correction between chips was executed using the Combat algorithm. The limma package facilitated the identification of differentially expressed genes between the control and disease groups, adhering to the criteria of P-value < 0.05 & |logFC| > 1. Furthermore, we downloaded the single-cell data of GSE157827 from the NCBI GEO database for subsequent examination.

### 2. GO and KEGG Functional Analysis

We employed the Metascape database (www.metascape.org) for annotation and visualization to elucidate the biological functions and signaling pathways implicated in the disease’s onset and progression. The differentially expressed genes underwent gene ontology (GO) analysis and Kyoto Encyclopedia of Genes and Genomes (KEGG) pathway analysis. A minimum overlap of ≥3 and a P-value of ≤0.01 were deemed statistically significant.

### 3. Feature Selection via Lasso Regression and SVM Algorithm

For the selection of disease diagnostic biomarkers, we applied Lasso regression and the SVM algorithm. The Lasso algorithm utilized the "glmnet" software package. Additionally, SVM-RFE, a machine learning technique based on support vector machines, was deployed to optimize variable selection by eliminating feature vectors generated by SVM. The "e1071" software package was used to establish a support vector machine model, further determining the diagnostic significance of these biomarkers in the disease.

### 4. Immune Cell Infiltration Analysis

The CIBERSORT method, a prominent tool for evaluating immune cell types in microenvironments, was adopted. Operating on the principle of support vector regression, this method conducts deconvolution analysis on the expression matrix of immune cell subpopulations. It includes 547 biomarkers and discriminates 22 human immune cell phenotypes, such as T cells, B cells, plasma cells, and myeloid cell subsets. In our study, the CIBERSORT algorithm was utilized to analyze patient data, estimating the relative proportions of 22 immune infiltrating cells. Additionally, Spearman correlation analysis was performed between gene expression levels and immune cell content.

### 5. GSEA Analysis

Gene Set Enrichment Analysis (GSEA) employs predefined gene sets to rank key genes based on their differential expression levels in two distinct sample types. It then determines whether these predefined gene sets are concentrated at the top or bottom of this ranking table. This study utilizes GSEA to compare signaling pathway differences between high-expression and low-expression groups, aiming to investigate the molecular mechanisms of core genes in both patient groups. The analysis is configured with 1000 permutations, with the permutation type set to phenotype.

### 6. Regulatory Network Analysis of Key Genes

In this research, the "RcisTarget" R package was employed for predicting transcription factors. RcisTarget calculations are motif-based, where the Normalized Enrichment Score (NES) of a motif is contingent on the total motifs in the database. Beyond the motifs annotated by source data, we extrapolated additional annotation files through motif similarity and gene sequence analysis. The initial step in assessing motif overexpression on a gene set involves calculating the Area Under the Curve (AUC) for each motif-gene set pair. This calculation is derived from the recovery curve of the motif ranking within the gene set. The NES for each motif is then determined based on the AUC distribution across all motifs in the gene set. The database "rcistarget.hg19.motifdb.cisbpont.500bp" was utilized for Gene-motif rankings.

### 7. cMap Drug Prediction

The Connectivity Map (CMap), developed by the Broad Institute, is a gene expression profile database centering on intervention gene expression. It is predominantly used to discern functional connections among small molecule compounds, genes, and disease states. The database encompasses gene chip data for 1309 small molecule drugs, including pre- and post-treatment scenarios in five human cell lines. Treatment conditions vary, encompassing diverse drugs, concentrations, and durations. This study predicts targeted therapeutic drugs for diseases through the analysis of differentially expressed genes.

### 8. Single Cell Analysis

Initially, the data was processed using the Seurat package, followed by an analysis of cluster positional relationships using the tSNE algorithm. Subsequently, clusters were annotated using the celldex package, focusing on cells significantly associated with disease onset. Lastly, marker genes for each cell subtype were extracted from the single-cell expression profile, setting the logfc.threshold parameter of FindAllMarkers to 1. We downloaded the single-cell datasets GSE129308 to study the mechanisms related to key genes in the new Braak VI AD prefrontal cortex dataset. Screen astrocytes from annotated data, use Seurat package, perform cell subpopulation analysis through dimensionality reduction clustering, perform cluster Profiler enrichment analysis on the marker genes of each subpopulation to determine the function of each subpopulation, then calculate the proportion of each subpopulation in the normal AD group, use AUCell for single-cell gene set scoring, genes from the Nrf2 pathway in the KEGG database, and plot ggplot2.

### 9. Statistical Analysis

Statistical analyses were executed using the R language (version 4.0). All tests were two-sided, with a p-value <0.05 deemed as statistically significant.

## Author contributions

Contribution to data analysis: W.C, L.Z, G.L.X. Contribution to data interpretation: W.C, R.F.Z, H.X.L. Contribution to writing the manuscript: W.C. All authors edited and approved the manuscript.

## Data availability

All data can be accessed through the National Center for Biotechnology Information (NCBI) Gene Expression Omnibus (GEO) database to access real-life data., Login numbers are GSE5281, GSE122063, GSE157827, GSE129308

## Animals and Western blot

Double transgenic mice App/Psen1(PS1/ΔE9) and Western blot were completed by Company from ShangHai Model Organisms. The protocol was approved by Experimental Animal Management and Use Committee of Shanghai Model Organisms Co., Ltd. The experiment involves the use of 5% w/v TBS extracts from brain homogenates were used for western blot analyses. The blot was probed with the following antibodies: anti-MAFF(Millipore); anti-MYL5(Millipore); anti-CDC37(Millipore). The band density was all normalized to β-actin. The intensities of the bands were measured using image J software.

## Authors

Wei Cui, Liang Zhang, Li Fang-Rui, Xi Huang, Gui-Lin Xie Northeast Agricultural University

