## Supplementary figures and images for "Combining machine learning algorithms and single-cell data to study the pathogenesis of Alzheimer’s disease"

### Supplementary document S1

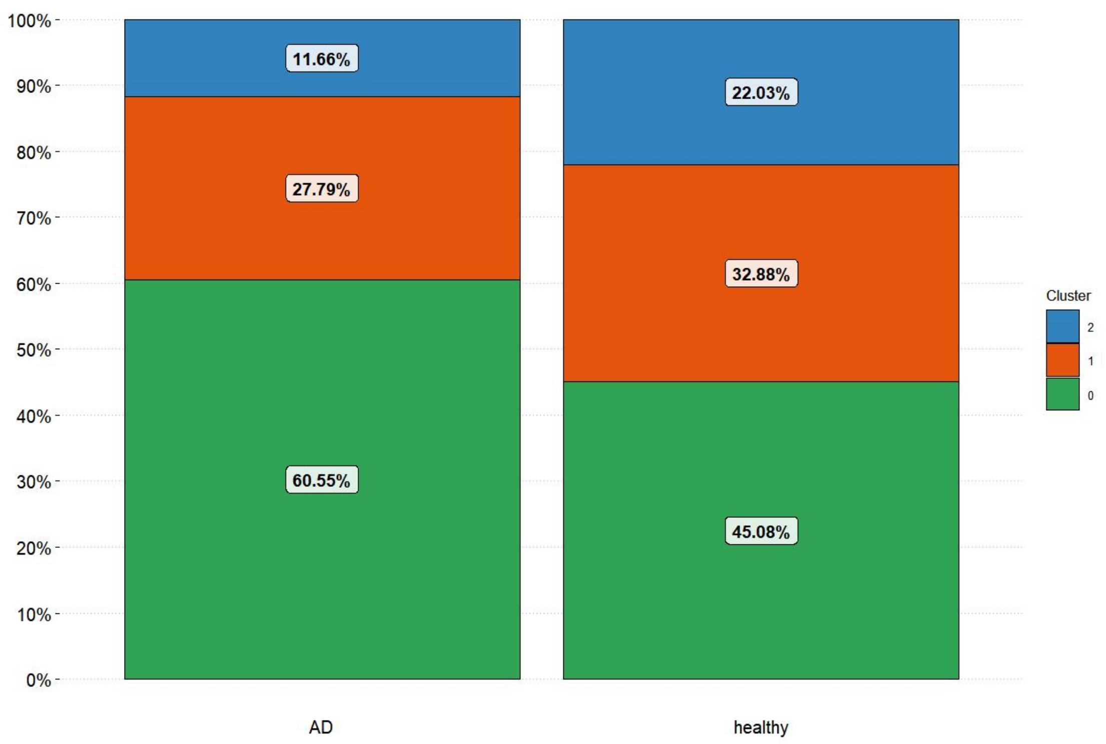

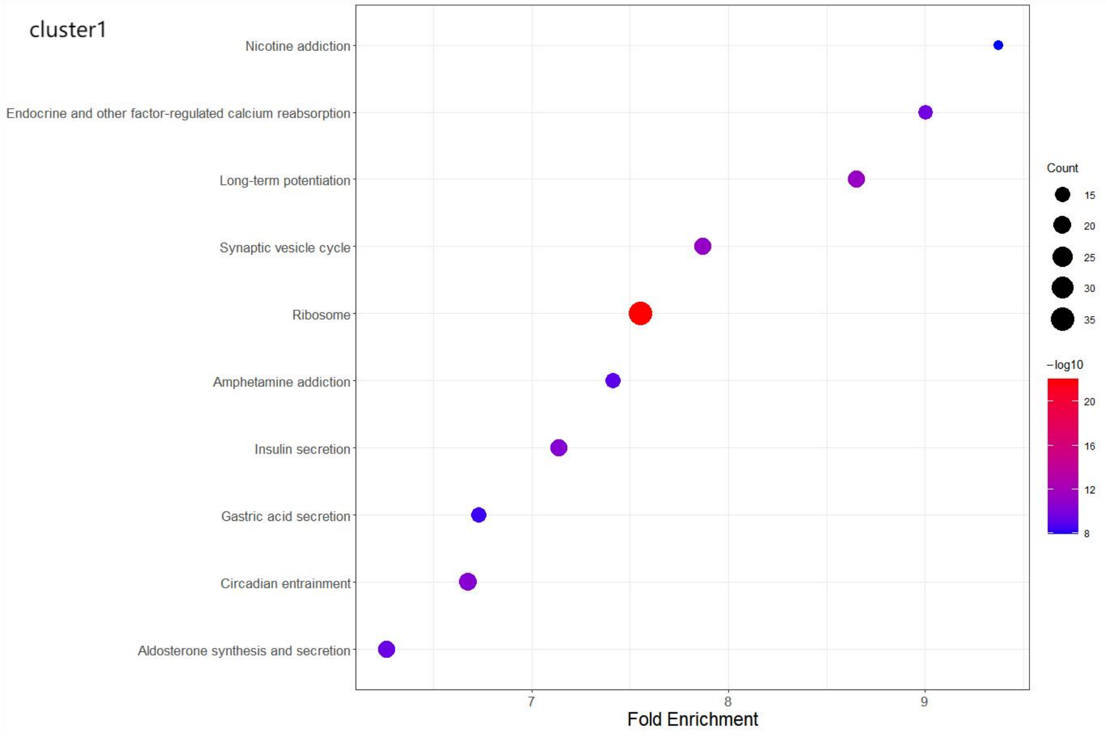

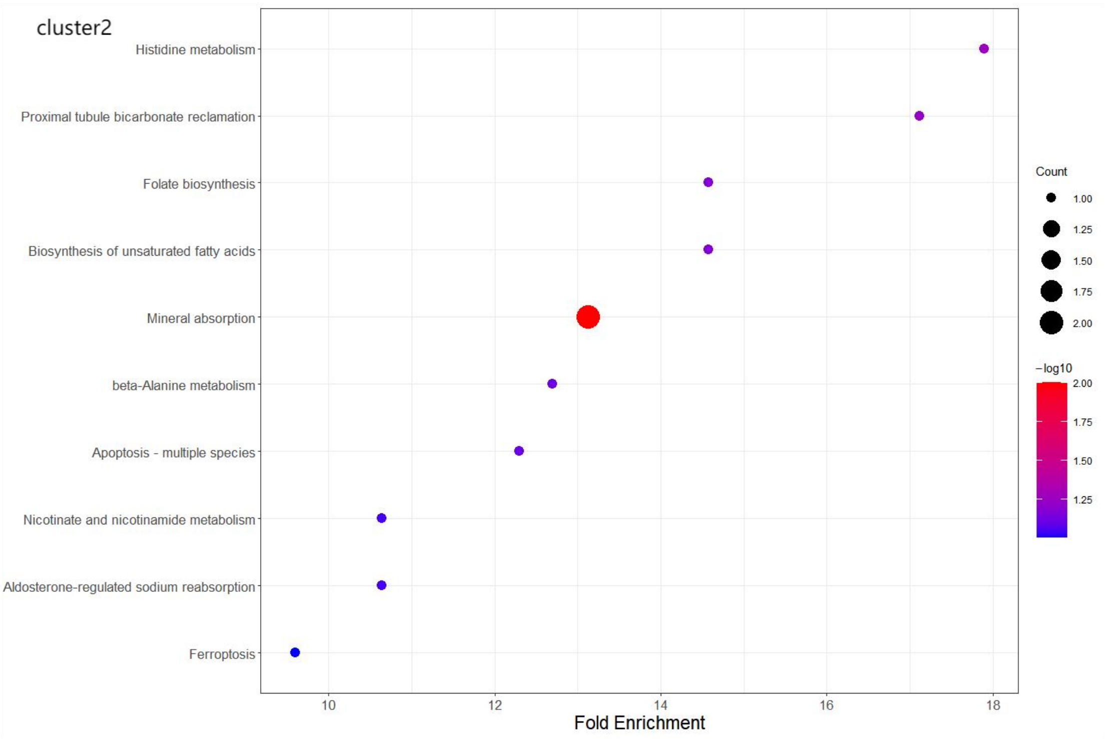

### Supplementary document S1

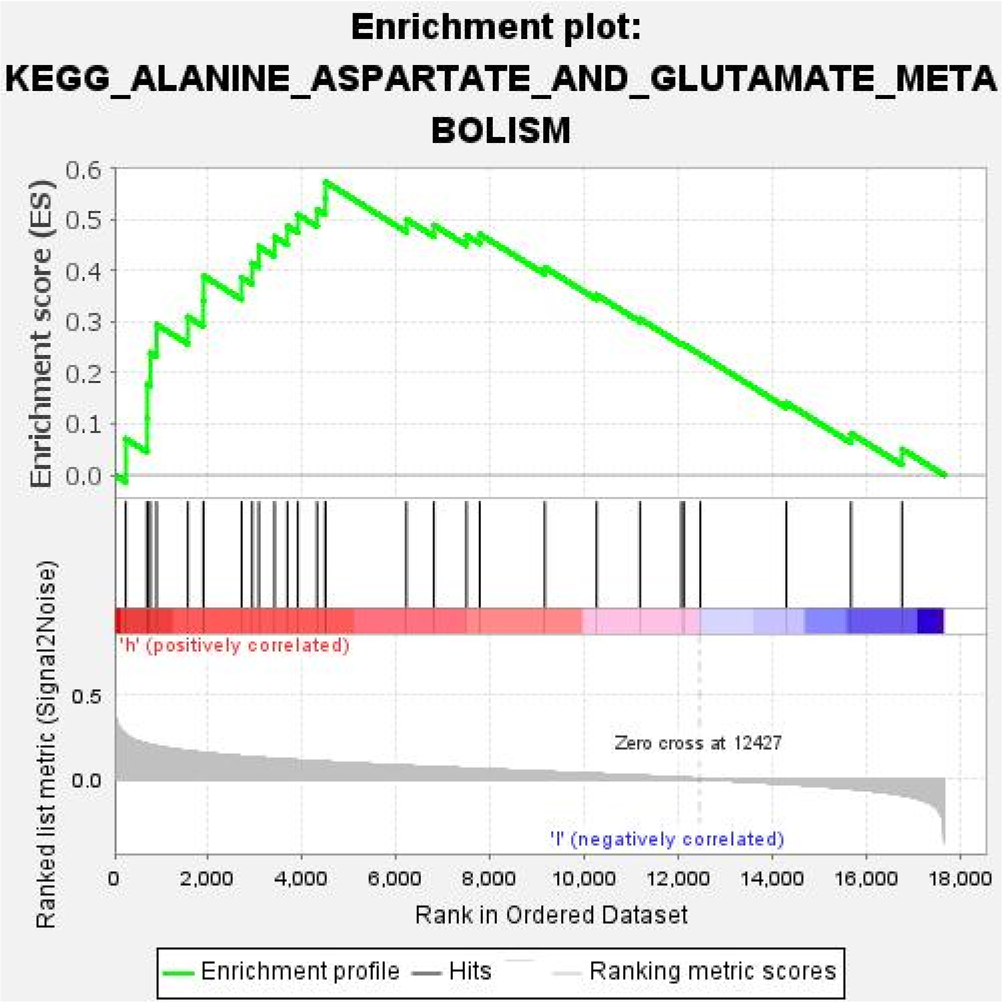

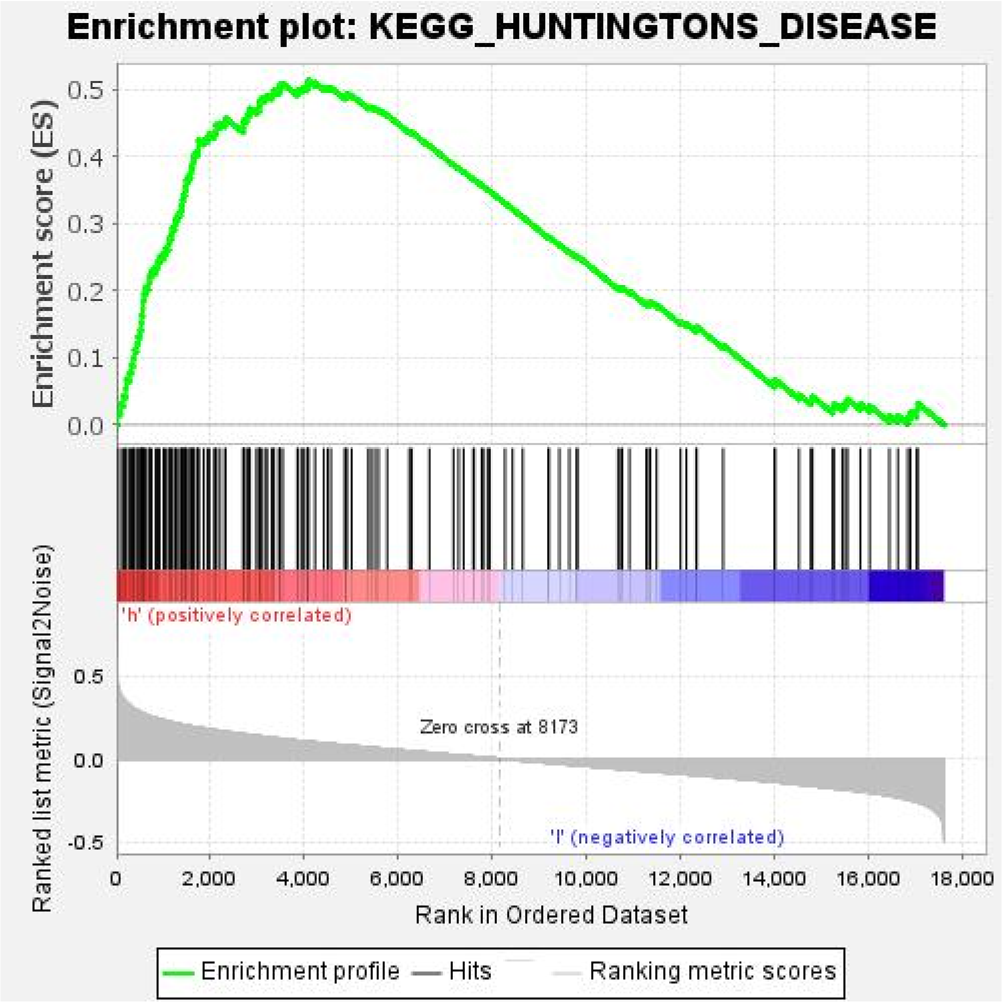

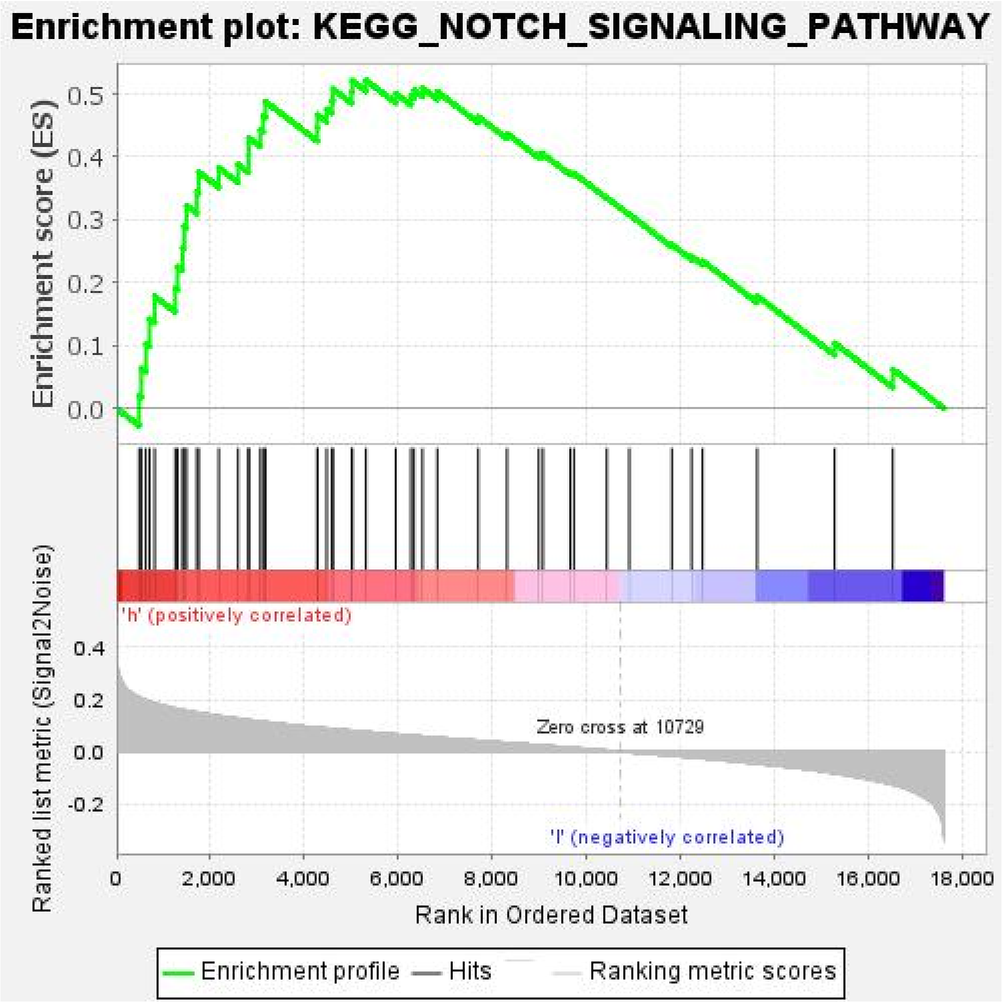

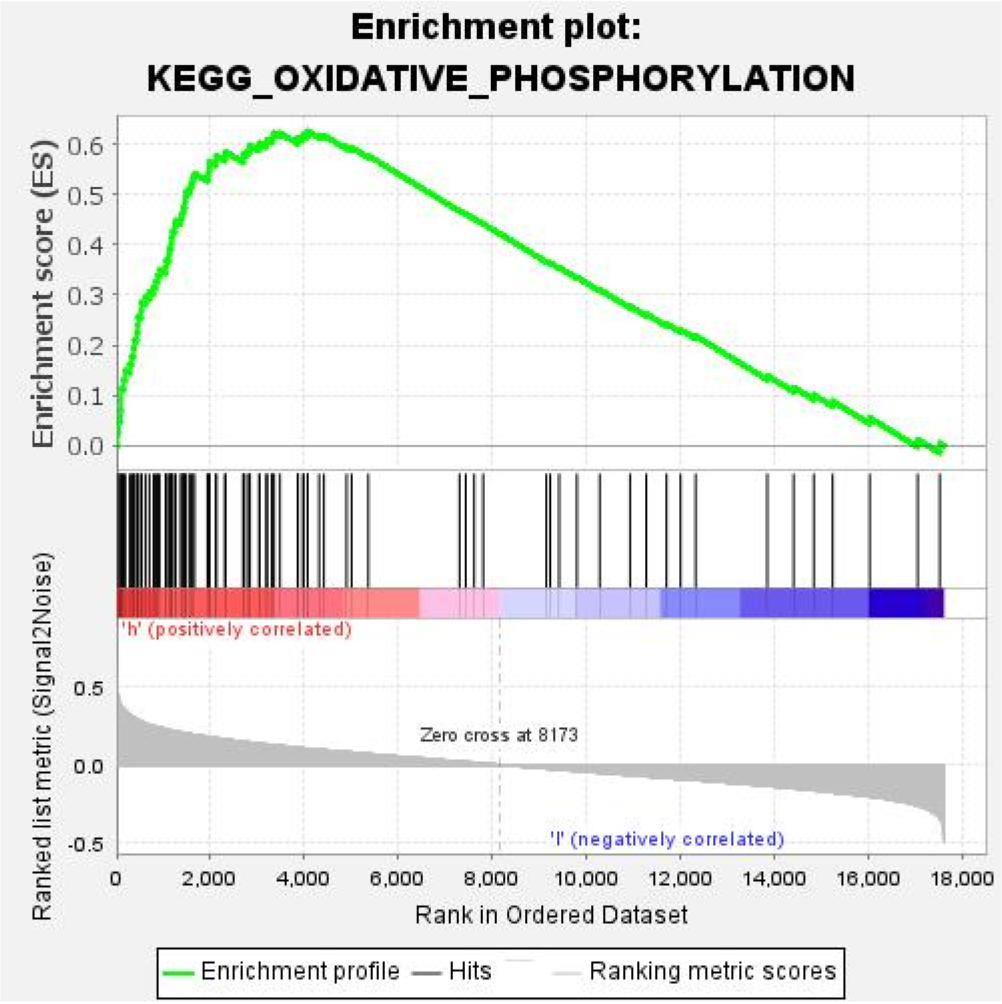

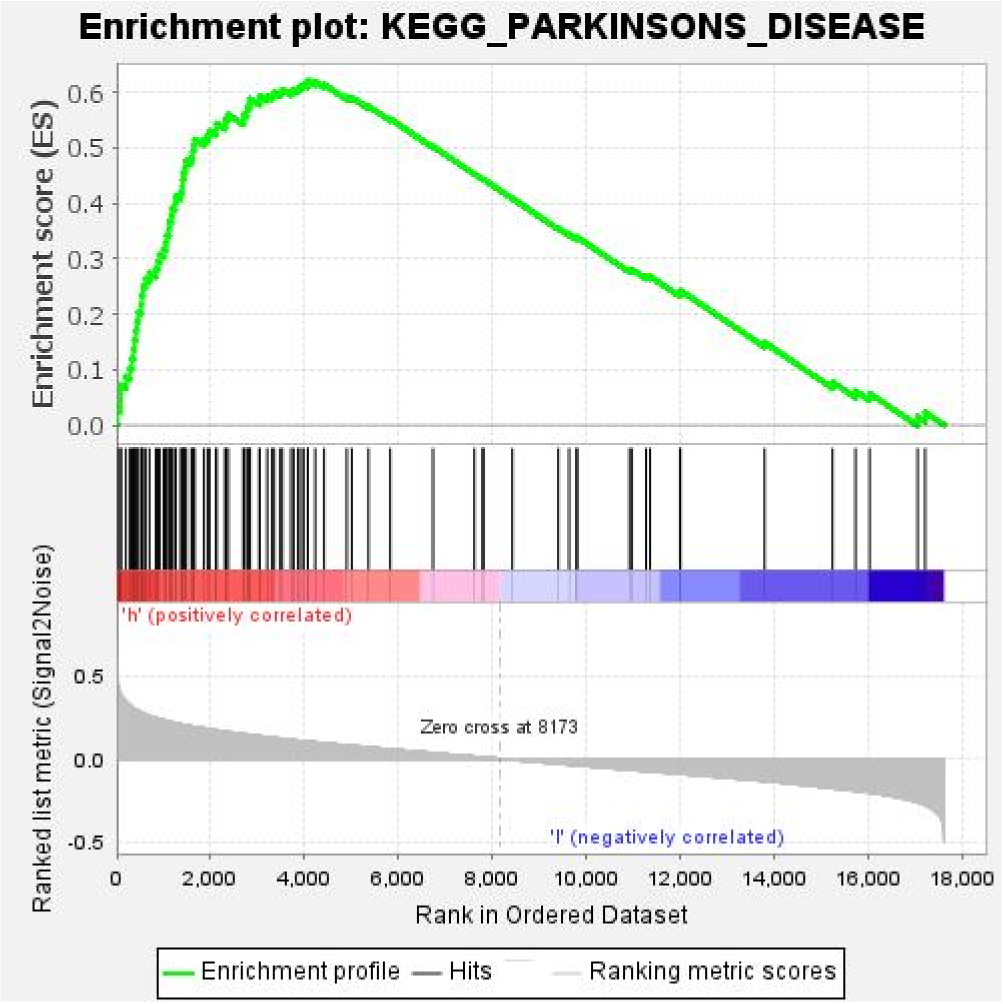

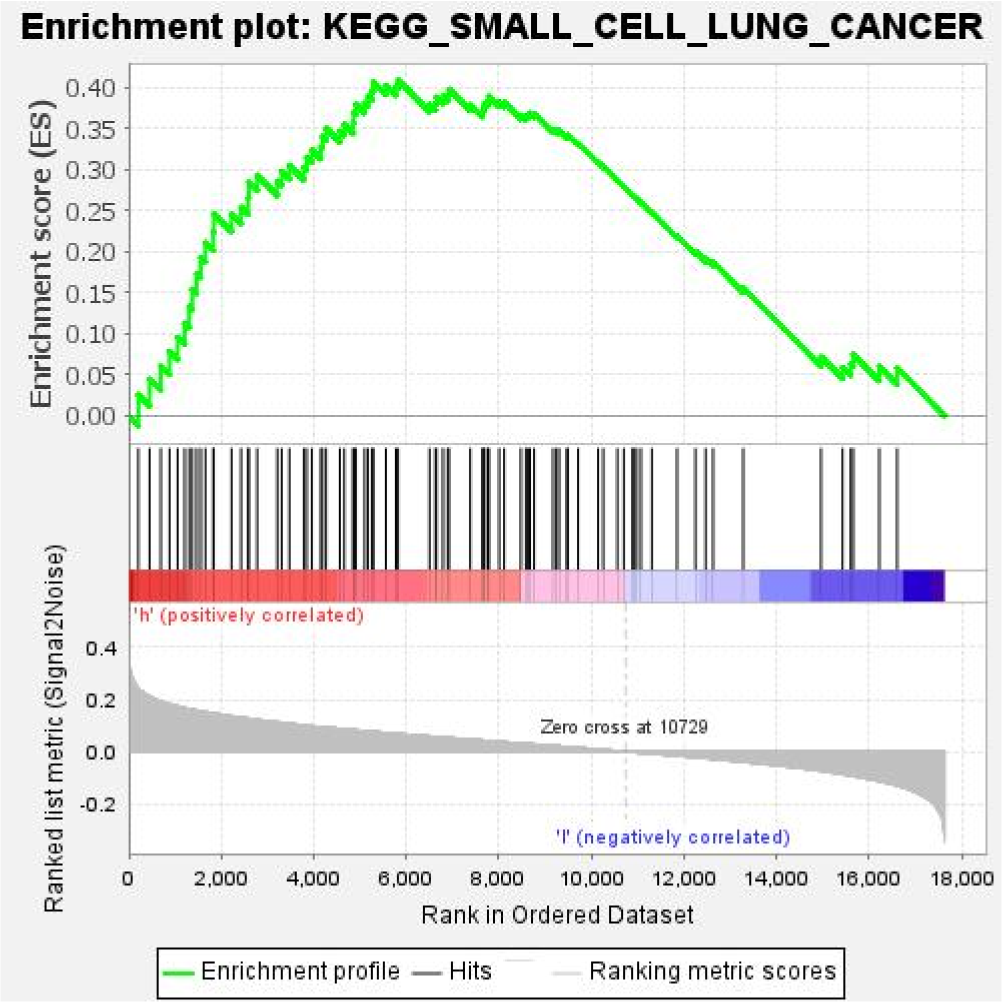
